# Endoderm Morphogenesis Reveals Integration of Distinct Processes in the Development and Evolution of Pharyngeal Arches

**DOI:** 10.1101/260356

**Authors:** Kazunori Okada, Hiroshi Wada, Shinji Takada

**Author notes:** Corresponding Authors: K.O.,; S.T.

## Abstract

The vertebrate pharyngeal arches (PAs) are established by a combination of two styles of segmentation; the most anterior 2 PAs are simultaneously but the others are sequentially formed. However, the mechanism underlying their coexistence is unclear. Here, we show that the simultaneous and sequential segmentation discretely proceeded, respectively, but were finally integrated at the second PP (PP2), by dynamic morphogenesis of pharyngeal endoderm in the zebrafish. The coordination of these 2 distinct processes appears to be common in the PA development of many vertebrates, in which specific developmental defects posterior to the PP2 are caused by mutations of particular genes or perturbation of retinoic acid signaling. Surprisingly, comparative analysis of PA segmentation showed that the combinatorial styles of PA development is present in shark but not in lamprey, suggesting that PA segmentation was modified in the stem gnathostomes corresponding to the drastic pharyngeal innovations, such as PA2-derived opercular.

## INTRODUCTION

The elaborated morphologies of organisms are often traced back to the simple metameric motifs, which are transiently established during development. Through the segmentation of these metameric motifs, the equivalent units, consisting of a certain group of cells, are formed and arranged along the body axis [1]. Each of these units becomes subsequently specialized to develop particular characteristics, based on their positional values, which are defined by collinear expression of Hox genes [2]. During vertebrate development, conspicuous segmental structures called pharyngeal arches (PAs) are bilaterally arranged in the ventral region of the head [3, 4]. PAs give rise to the segmental organization of skeletons, muscles, nerves, and vessels in the pharynx; and, therefore, segmentation and subsequent specification of the PA are crucial for the development of the vertebrate head [5].

In addition to all 3 germ layers, PA development involves cranial neural crest cells (CNCCs) [3-5]. Importantly, CNCCs have been considered to be dominant in the differentiation of PA-specific characteristics [6, 7]. The tripartite streams of CNCCs, referred to as the trigeminal, hyoid, and branchial streams, specify the regions of the jaw-forming first PA (PA1) or mandibular arch (MA); the hyoid-forming PA2, called the hyoid arch (HA); and the more posterior PAs, referred to as the branchial arches (BAs), respectively [8-11], depending on branchial Hox codes specified in the CNCCs [12]. On the other hand, the segmentation of the PA units occurs independently of these CNCCs [13, 14]. Rather, the pharyngeal endoderm plays a pivotal role in this segmentation by generating epithelial outpocketings called pharyngeal pouches (PPs), which physically define the anterior and posterior interfaces of each PA [4, 5].

Interestingly, the anterior most 2 PAs (MA and HA) are formed simultaneously; whereas the posterior PAs (BAs) are generated sequentially in an anterior to posterior order [13, 15, 16], suggesting discrete regulation of the anterior and posterior PA segmentation. Correspondingly, retinoic-acid (RA) deficiency in zebrafish [17], quail [18], rat [19] and mouse [20] embryos consistently results in abnormalities in the segmentation of their PAs posterior to the second PP (PP2). Similarly, *pax1* -knockout in medaka [21] and *Ripply3*-knockout in the mouse [22] show the loss of the posterior PAs but a normal MA and HA. These studies suggest that distinct segmentation mechanisms for the anterior and posterior PAs may operate to establish the entire series of PAs. However, the development of this complex style of PA segmentation has been poorly understood. In particular, the question as to how the two distinct mechanisms are integrated at the formation of the PP2 should be answered.

This unique segmentation of PAs probably was established during the course of vertebrate evolution. The vertebrate PPs are homologous to the endodermal gill slits in the pharynx of non-vertebrate deuterostomes, such as hemichordate and amphioxus [4, 23-26]. In contrast to the segmentation of vertebrate PPs, all of these gill slits are simply formed in a sequential manner [24, 27]. Since similar sequential segmentation occurs only in the BA region of vertebrates, the endodermal segmentation had probably been modified in the evolution of the vertebrate lineage. In addition to the evolution of the segmentation style, innovative roles of the endoderm in pharyngeal development should also have arisen in accordance with the acquisition of neural crest cells by the common ancestor of vertebrates. Previous studies have shown that not only the intrinsic machinery in CNCCs but also signals from the PP endoderm to the CNCCs are crucial for the development of the cranial skeletons [16, 28-30]. Therefore, it is plausible to propose that the pharyngeal endoderm experienced some developmental modifications that enabled it to regulate the development of the CNCC-derived skeletons during vertebrate evolution [4]. However, the evolutionary scenario of PP segmentation and its contribution to the innovations of the vertebrate pharyngeal apparatus remain to be elucidated, mainly owing to the lack of adequate understanding of the mechanism of PP development.

Recent studies on zebrafish PP development have revealed the dynamic cellular nature of the endoderm forming the PPs [31-34], showing the advantages of the zebrafish model to dissect the processes of PP development. In this study, we examined the development of the zebrafish pharyngeal endoderm, especially focusing on the formation of PP2. Precise examination by live-imaging and cell-tracing experiments performed in zebrafish showed that the morphogenesis of the anterior and posterior PAs were apparently distinct. Especially, we found that PP2 was formed in an unexpected manner; i.e., the rostral and caudal aspects of PP2 were initially formed separately, then subsequently accessed with each other by dynamic remodeling of endoderm epithelium, and finally became integrated. These results resolved the pending issue regarding the interface between the distinct mechanisms of PA development. Crucially, this style of PA development was never established in the lamprey, an extent jawless vertebrate; whereas it became shared among the gnathostomes (jawed vertebrates). Thus, our findings also suggested that renovation of PA segmentation in the gnathostome lineage likely contributed to the evolution of the vertebrate craniofacial skeletons.

## RESULTS

### Rostral and Caudal Aspects of PP2 Emerged Separately during the Development of Zebrafish Pharyngeal Endoderm

To better understand the development of the PA endoderm, we performed time-lapse imaging of the endodermal cells in transgenic zebrafish *Tg(sox17:EGFP)*, in which EGFP expression was specifically driven in the endodermal cells by the *sox17* promoter [35]. PA1 and PA2 appeared simultaneously at 16 hours post fertilization (hpf; Figure 1A-1F, Movie S1), whereas PP outpocketings posterior to PP3 were sequentially generated in an anterior-to-posterior order after 16 hpf (Figure 1F-1J, Movie S2). Unexpectedly, we found that 2 endodermal bulges appeared simultaneously with the PP1 budding in the area where the PP2 would be generated (Figure 1C, Movie S1). Notably, these bulges were gradually remodeled and finally became integrated with each other to form the PP2 (Figure 1G-1N, Movie S2). This remodeling occurred not prior to but in parallel with the sequential generation of the posterior PPs, suggesting that the posterior development proceeded independent of that of the PP2 integration (Figure 1G-1N, Movie S2).

**Figure 1.**
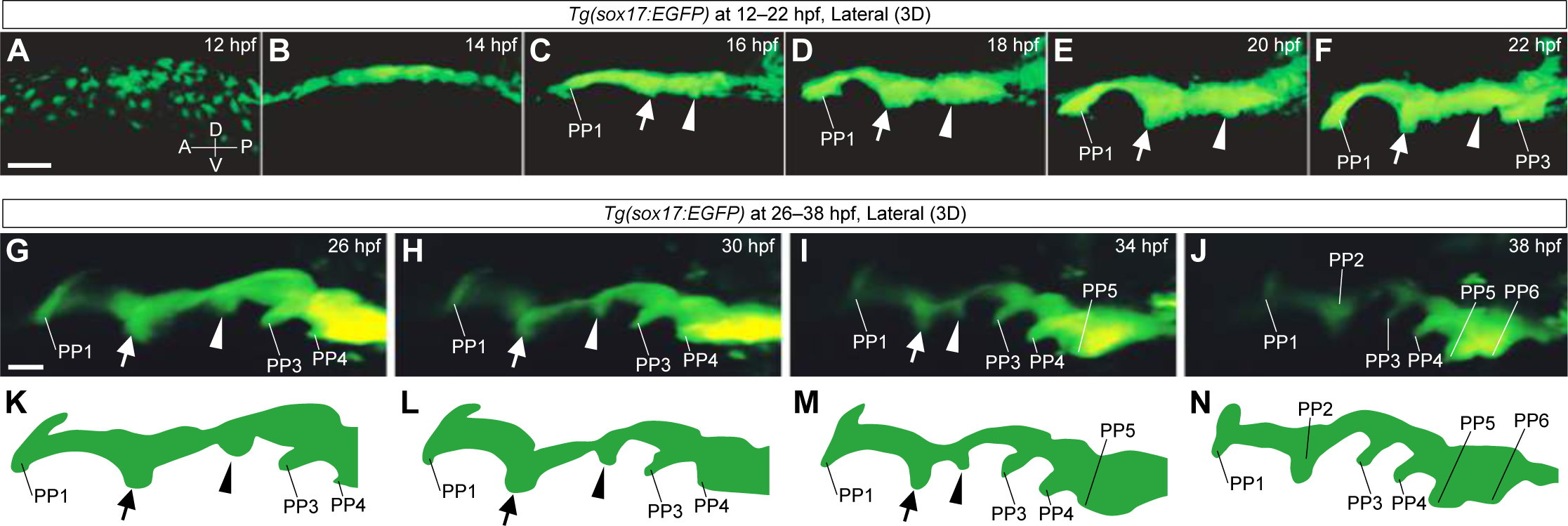
Time-lapse observations of the pharyngeal endoderm during PP segmentation in *Tg(sox17:EGFP)* zebrafish embryos. (A-J), Time-lapse analysis of the pharyngeal endoderm of *Tg(sox17:EGFP)* zebrafish from 12 to 22 hpf (A-F, Movie S1) and from 26 to 38 hpf (G-J, Movie S2). Rostral (arrow) and caudal (arrowhead) bulges appeared posterior to PP1 and gradually fused to form a PP2. (K-N), Schematic illustrations of the shape of the lateral pharyngeal endoderm in G-J, respectively. A, anterior; P, posterior; D, dorsal; V, ventral; PP1–6, the first to sixth pharyngeal pouches; arrows, rostral bulges; arrowheads, caudal bulges. Scale bars, 50 μm (A) and 20 μm (G).

To understand the dynamism of the endodermal bulges, we performed a lineage-tracing experiment by means of endoderm-specific photoconversion. To this end, we created *Tg(sox17:Kaede)*, a transgenic line harboring *kaede* expression under the control of the *sox17* promoter. Photoconversion of cells in the rostral bulge at 20 hpf revealed that these cells contributed to the rostral aspect of PP2 at 48 hpf (Figure 2A-2E and Figure S1). The descendants of these cells extensively spread to form the inner lining of the distal part of the HA, where an opercular flap would later expand (Figure 2A-2E and Figure S1). On the contrary, descendants from the caudal bulge became distributed in the caudal aspect of PP2 at 48 hpf, especially to its proximal region (Figure 2F-2J and Figure S1). In addition, cells in the intermediate region between the rostral and caudal bulges contributed to the more distal and ventral regions of the caudal aspect and the dorsal edge of the PP2 (Figure 2K-2N and Figure S1). Based on the various patterns of cell traces (n = 29, Figure 2 and Figure S1), we obtained an overview of the endodermal cell fate in the future PP2 region at 20 hpf (Figure 2O). Thus, the PP2 was generated by the dynamic remodeling of endodermal cells between the rostral and the caudal bulges, which directly contributed to the respective rostral and caudal aspects of PP2.

**Figure 2.**
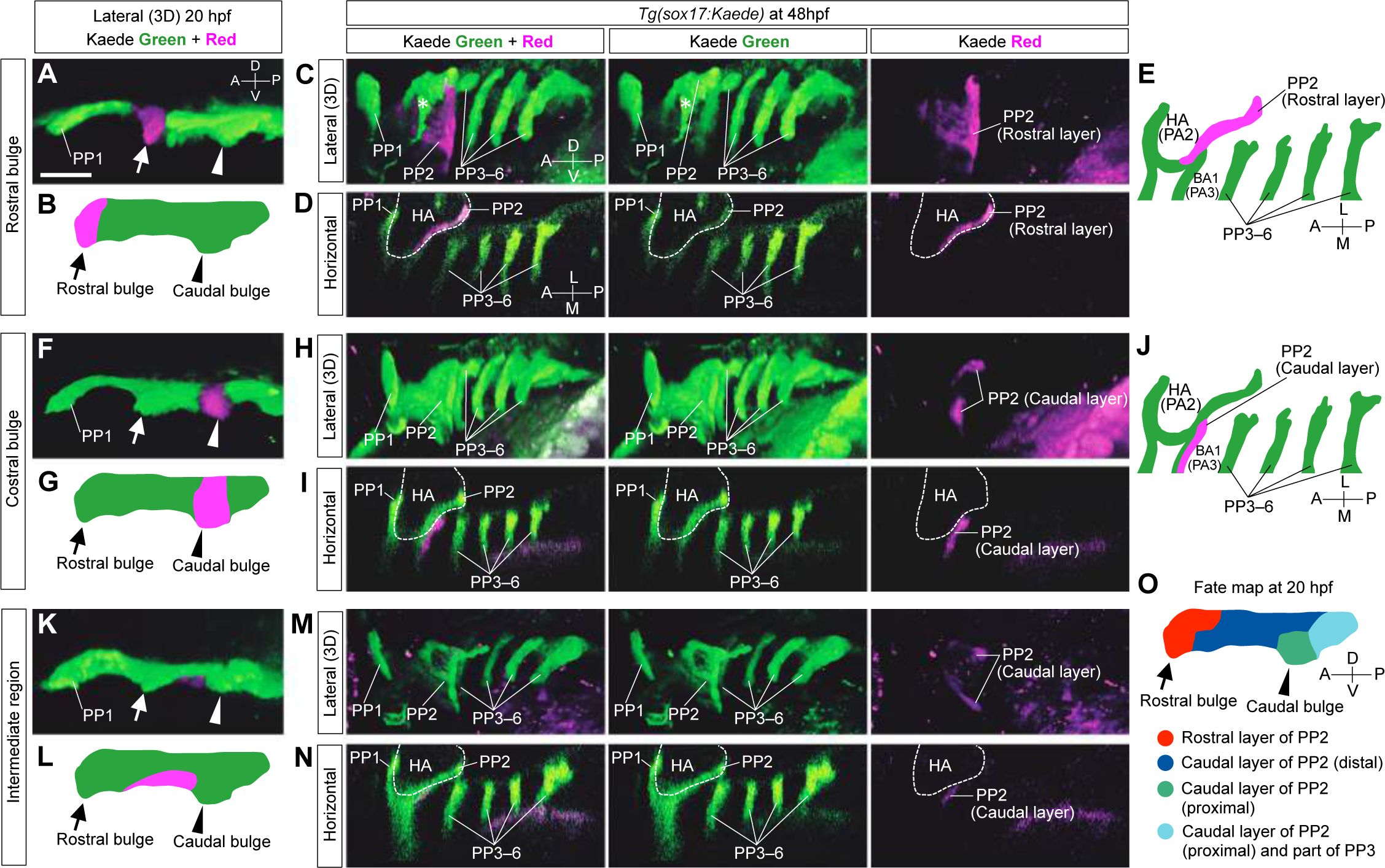
Lineage tracing of endodermal cells in *Tg(sox17:Kaede)* zebrafish embryos by photoconversion. (A-E) The cells of the rostral bulge (arrows) were marked at 20 hpf (A and B). At 48 hpf, cells of the rostral bulge contributed to the large area of the rostral aspect of PP2 (C-E). (F-J) The cells of the caudal bulge (arrowheads) were marked at 20 hpf (F and G). At 48 hpf, the descendant cells contributed to the caudal aspect of PP2 rather proximally (H-J). (K-N) The cells of the intermediate domain of a putative PP2 (between the rostral and caudal bulges) were marked (K and L). At 48 hpf, these descendants composed the dorsally and ventrally distant area in the caudal aspect of PP2 (M and N). (O) Overview of the cell fate in the future PP2 endoderm at 20 hpf. Cell fates were examined in various regions of the presumptive PP2 endoderm by photoconversion (n = 26, A-N and Figure S1); and these are summarized, showing the dynamic reorganization of the endoderm forming PP2. A, anterior; P, posterior; D, dorsal; V, ventral; M, medial; L, lateral; BA, branchial arch; HA, hyoid arch; PP1–6, the first to sixth pharyngeal pouches; arrows, rostral bulge; arrowheads, caudal bulge; asterisk, blood vessel. Scale bar, 50 μm.

### Rostral and Caudal Bulges of Future PP2 Endoderm Region Exhibited Characteristics of the Pharyngeal Pouch

Our time-lapse observations and cell-tracing experiments revealed that PP2 arose from 2 independent endodermal domains, which then coalesced into a pouch structure by the dynamic remodeling of the endoderm. To understand the development of PP2 more precisely, we examined the expression of PP-specific genes in the future PP2 endoderm. Remarkably, expression of *nkx2.3*, which is observed in the zebrafish PPs [36], was specifically detectable in the rostral and caudal bulges at 20 hpf (Figure 3A and Figure S2). Consistent with the PP2 maturation, these *nkx*2.3-positive bulges gradually converged to form PP2 (Figure S2). Thus, PP2 was separately established prior to the epithelial remodeling at the rostral and caudal bulges. We also examined the expression of *pax1*, which is known to be expressed in the PPs of many vertebrates. In the zebrafish genome, there are 2 orthologs of *pax1, pax1a* and *pax1b*. As expected, *pax1b* was expressed in the rostral and caudal bulges, as well as in the other PPs (Figure S2). On the other hand, *pax1a* was expressed in the caudal, but not the rostral bulge, suggesting that the rostral bulge might exhibit some specific character different from that of the other PP endoderm (Figure 3E-3H). These expression patterns of the PP-specific genes support the idea that at least some characteristics of PP2 had already been provided in the bulges. Furthermore, the separate expression of *nkx2.3* in the 2 distinct bulges support our results obtained by time-lapse observation and cell-tracing experiments, which showed that PP2 was formed not by simple bending of the future PP2 region but by complex remodeling of the 2 distinct endodermal bulges.

**Figure 3.**
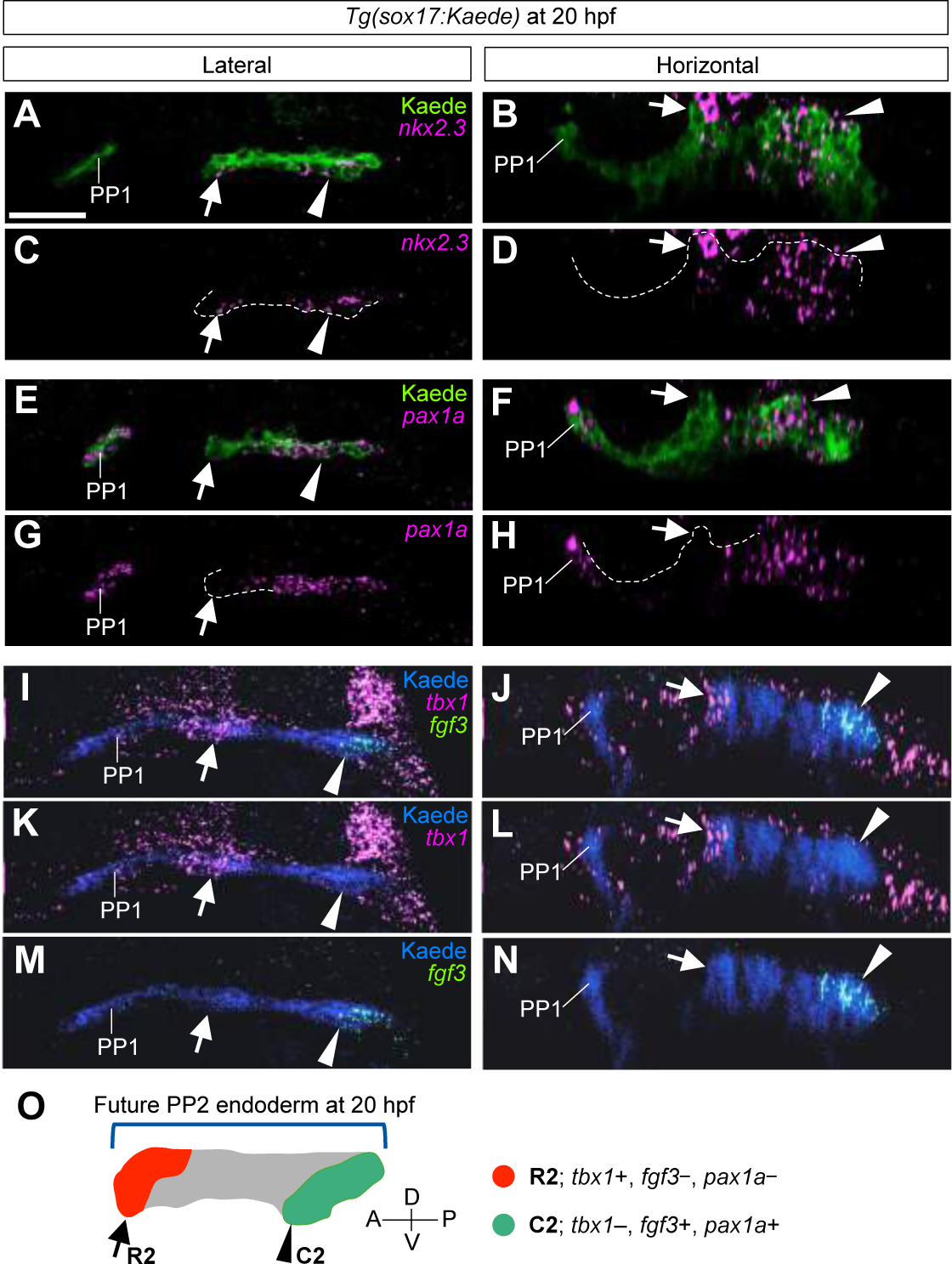
Separated formations and the rostrocaudal identity in the PP2 endoderm at 20 hpf. (A-D) Expressions of *nkx2.3* separately indicated the rostral (arrows) and caudal (arrowheads) bulges of the PP2 endoderm indicated by immunohistochemistry with Kaede antibody. (E-H) Expression of *pax1* was detected in PP1 and the caudal part of PP2 endoderm (arrowheads) but almost absent in the rostral bulge of the PP2 endoderm (arrows). (I-N) Expressions of *tbx1* (I-L) and *fgf3* (I, J, M and N) were detected in the respective regions of rostral (arrows) and caudal (arrowheads) bulges of PP2 showing an early specification of rostrocaudal polarity of the PP2. (O) According to the fate analysis and the molecular profiles, the rostral and the caudal bulges of the future PP2 are distinctly defined as R2 (red, arrow) and C2 (green, arrowhead), respectively. Whereas cells of the intermediate region (gray) contribute to the caudal aspect, *fgf3*, which is a caudal marker, is not expressed in these cells at 20 hpf. A, anterior; P, posterior; D, dorsal; V, ventral; PP1, the first pharyngeal pouch; arrows, rostral bulge; arrowheads, caudal bulge. Scale bar, 20 μm.

To further understand the development of the future PP2 endoderm, we examined the rostro- and caudal-specific molecular characteristics of the PPs. The expression of *tbx1*, which was specific to the rostral aspect of each PP (Figure S2), was strongly detected in the rostral bulge but not in the caudal bulge (Figure 3I-3L); whereas that of *fgf3*, specific to the caudal aspect of PPs (Figure S2), was detected in the caudal bulge but not in the rostral one (Figure 3l, 3j, 3M and 3N). Crucially, *tbx1-* and fgf3-positive domains were separately detected in the PP2 endoderm before its integration, indicating that the rostral and caudal bulges had already acquired distinct rostrocaudal characteristics prior to the epithelial remodeling to form PP2. We refer to these rostral and caudal bulges as R2 and C2, respectively, hereinafter (Figure 3O).

### R2 and C2 Independently Contributed to Skeletal Development in HA and BA

Since the molecular characteristics of the rostral and caudal aspects of PP2 were observed in the endodermal domains of R2 and C2, respectively, we next investigated whether the rostral and caudal identities had actually been determined in R2 and C2. To this end, we specifically ablated the EGFP-labeled endodermal cells in the R2 or C2 region of *Tg(sox17:EGFP)* embryos by infrared laser-mediated heating [37, 38].

Consistent with the results from the cell-lineage tracing, ablation of R2 cells at 20 hpf (Figure 4A and 4A’) impaired the expansion of the rostral aspect of PP2 at 48 hpf (n = 3/3, Figure 4B and 4C). In later stage, R2 ablation resulted in loss of HA-derived dermal bones of the branchiostegal ray (BR; n = 12/16) and the opercular (OP; n = 10/16), both of which compose the operculum (Figure 4D-4F). For operculum development, Shh is required to be expressed in the HA [39]. We found that *shha* was expressed in the endoderm corresponding to the R2-derived cells (Figure 4G and 4H) and that this expression was decreased (n = 5/12) or eliminated (n = 5/12) by R2 ablation (Figure 4I), suggesting that the R2 region gave rise to a signaling center of Shh for operculum formation. In addition, this ablation occasionally caused a size reduction of other HA-derived skeletons, such as the hyomandibular (HM; n = 3/16) and ceratohyal (CH; n = 6/16). The ceratobranchial (CB) cartilages, however, which are derived from BAs, were completely normal (n = 16/16, Figure 4D).

**Figure 4.**
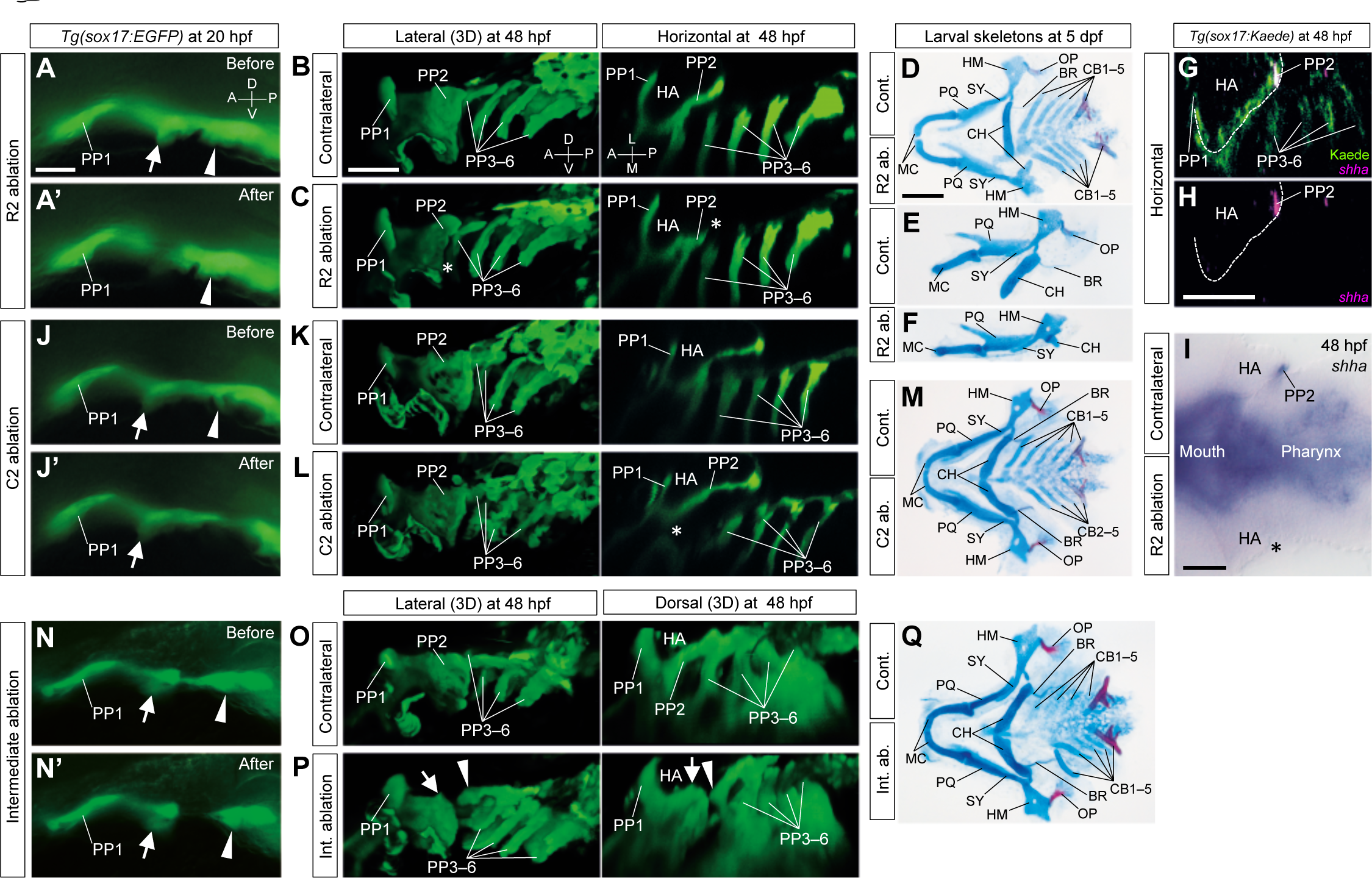
Early determinations of distinct roles for later skeletal patterns in R2 and C2 endoderm. (A and A’) Cells of R2 (arrows) in *Tg(sox17:EGFP)* embryos were ablated at 20 hpf. (B-E) R2 ablations caused a specific loss of the epithelial expansion of the caudal lining of HA (asterisks in C) and reductions in HA-derived skeletons, especially in the opercular series (OP, BR) (D-F). (G and H) Expression of *shha*, required for opercular development, was detected in the PP2 endoderm occupied by the R2 descendants (G and H). (I)Consistent with the endodermal (B and C) and the skeletal (D-F) phenotypes, R2 ablation caused a specific loss of the *shha* expression in PP2, as shown in a flat-mounted embryo (asterisk in I). (J and J’) Cells of C2 (arrowheads) in *Tg(sox17:EGFP)* embryos were ablated at 20 hpf. (K-M) Ablations of C2 cells caused a loss of the proximal region of PP2, which consists of the rostral lining of the third PA (BA1) *(K* and L, asterisk in L), resulting in a loss of CB1 cartilage (M). (N and N’) Endodermal cells between R2 (arrows) and C2 (arrowheads) were ablated in *Tg(sox17:EGFP)* embryos at 20hpf. (O-Q) Ablations of cells in the intermediate region did not affect the segregations of HA and BA1 but caused abnormal arrangements of them, shown by a split between HA and BA1 (O and P). Correspondingly, in the ablated sides, the positions of the BA1-derived CB1 cartilage shifted posteriorly; although a complete set of the pharyngeal skeletons developed (Q). Images of ablation sides (C, F, L and Q) were inverted in a left-right direction for comparisons with contralateral sides. A, anterior; P, posterior; D, dorsal; V, ventral; L, lateral; M, medial; BR, branchiostegal ray; CB1–5, the first to fifth ceratobranchials; CH, ceratohyal; HA, hyoid arch; HM, hyomandibular; MC, Meckel’s cartilage; OP, opercular bone; PP1–6, the first to sixth pharyngeal pouches; PQ, palatoquadrate; SY, symplectic; arrows, R2; arrowheads, C2. Scale bars, 20 μm (A), 50 μm (B, H and I) and μm (D).

In contrast, ablation of cells in the C2 region (Figure 4J and 4J’) caused abnormalities in the proximo-caudal PP2 adjacent to BA1 (or PA3; n = 3/3, Figure 4K and 4L), resulting in specific loss of the first CB cartilage (CB1; n = 8/8, Figure 4M). On the other hand, ablation of the intermediate cells between R2 and C2 (Figure 4N and 4N’) did not cause any loss of the pharyngeal skeleton; although the position of CB1 on the ablation side shifted posteriorly and laterally (n = 4/6, Figure 4Q). Interestingly, this ablation caused a split of endoderm between HA and BA1 (n = 10/12, Fig. 4 *O* and *P*), which were almost normally formed, showing the necessary role of the intermediate endoderm for the integration of HA and BA1. On the other hand, we could exclude the possibility that the loss of skeletal elements was due to possible deficits of CNCCs caused by the infrared irradiation to the endodermal cells, because the PA mesenchymal cells and expression of *dlx2a*, which is a credible marker of CNCCs in PAs, were not obviously changed by the ablation of the adjacent endoderm (n = 6, Figure S3). Therefore, we concluded that the endodermal cells of R2, C2, and the intermediate region played distinct roles for the craniofacial development in zebrafish. Significantly, their distinct roles were assigned or determined in the endodermal domains prior to the remodeling for the morphological maturation of PP2.

### Distinct Molecular Machineries for Rostral and Caudal Development of PP2: Mechanistic Interface and Integration Process of PA Development Unveiled

Previous studies suggested that the development of the anterior and posterior PAs appears to be distinct in the molecular mechanism [17-22]. These studies suggested that the mechanistic boundary between them exists in the formation of the PP2. Thus, to resolve the complex system of the PA development, understanding of the molecular mechanism of the PP2 development should be considered crucial. Since our findings enabled us to dissect the developmental process of the PP2, we next addressed the issue as to how the distinct molecular machineries could coordinately achieve PP2 and the series of the vertebrate PAs.

Since RA signaling is specifically required for the development of the posterior PPs [17-20], we supposed that the R2-C2 boundary would correspond to the anterior border of RA function. Visualization of RA activity by utilizing a transgenic fish *Tg(RARE:Venus)*, which harbors a Venus reporter driven by RA-responsive elements (RARE) [40], showed that Venus was expressed in the C2 cells at 20 hpf and further persisted in their descendants, as well as in the posterior pharyngeal endoderm (Figure 5A-5F). In contrast, Venus expression was never detected in the R2 and PP1 endoderm throughout our examination (Figure 5A-5F). Furthermore, treatment with DEAB, which inhibits RA biosynthesis, impaired C2 formation and *fgf3* expression in the C2 (Figure 5K-5N), but not *tbx1* expression in the R2 (Figure 5G-5J). Thus, RA signaling was specifically activated in and required for the pharyngeal endoderm posterior to the R2, indicating that the anterior border of RA function actually corresponded to the R2-C2 border we had identified.

**Figure 5.**
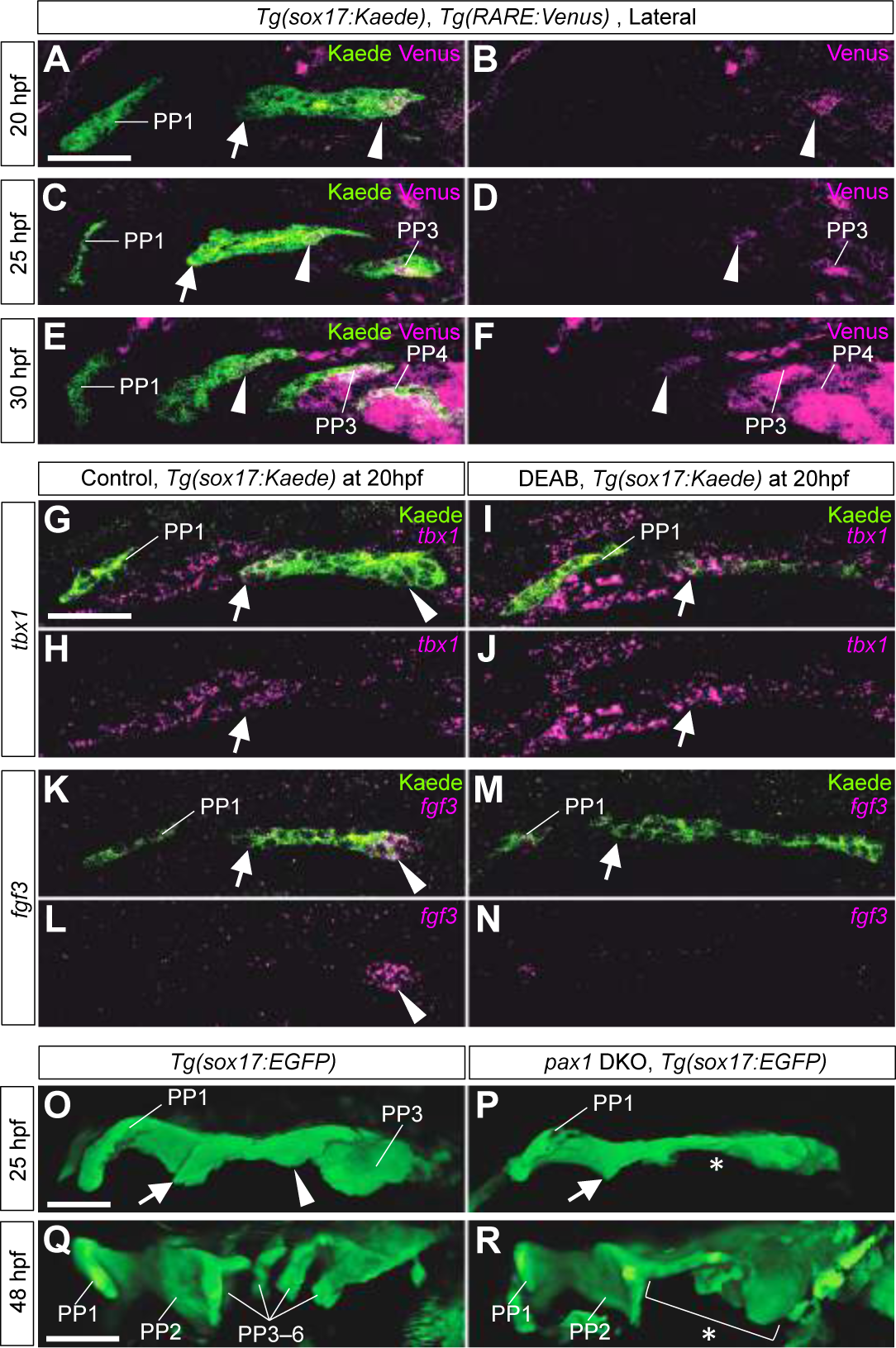
Boundaries of molecular mechanisms forming PPs between the rostral and caudal aspects of PP2. (A-F) Immunohistochemistry of double transgenic embryos of *Tg(sox17:Kaede)* and *Tg(RARE:Venus)* showed the specific signals of RA reporter Venus in the caudal aspect of PP2 and in the posterior PPs but not in the rostral aspect of PP2 and PP1 endoderm at 20 hpf (A and B), 25 hpf (C and D) and 30 hpf (E and F). (G-J) Expression of *tbx1* in R2 (arrows) was not affected. (K-N) RA deficiency caused by DEAB treatment resulted in a loss of *fgf3* expression in C2 (arrowheads). (O-R) Endodermal morphologies of wild-type (O and Q) and *pax1a;* pax1b-double knockout *(pax1* DKO) embryos (P and R) harboring a *Tg(sox17:EGFP)* transgene. At 25 hpf, PP1, R2, C2, and PP3 were formed in the wild type (O); but in the *pax1* DKO embryos, C2 and PP3 were specifically defective (P, asterisk). At 48 hpf, complete segments of PP were observed in the wild type (Q); whereas the caudal PP2 and more posterior PPs were not formed in *pax1* DKO embryos (R, bracket and asterisk). Notably, PP1 and the rostral aspect of PP2 were almost normal in the mutants (*R*). All pictures show the left side view of the pharyngeal region. PP1–6, the first to sixth pharyngeal pouches; arrows, R2; arrowheads, C2. Scale bars, 50 μm.

Our previous study also showed that *pax1* is specifically required for the development of posterior PPs in medaka [21]. Since zebrafish has 2 *pax1* homologs, *pax1a* and *pax1b*, we generated double knockout mutants of these genes *(pax1* DKO) by performing CRISPR/Cas9-mediated mutagenesis (Figure S4). As expected, the *pax1* DKO embryos clearly showed abnormalities in the development of their pharyngeal pouches posterior to the C2, but not in the R2 (Figure 5O-5R). Consistently, the gill skeletons, but not opercular skeletons, were lost in the *pax1* DKO larvae (Figure S4). Although the anterior part of HM (aHM), in which PP1 is required for its development [29], was lost in the *pax1* DKO larvae (Figure S4), the PP1 was normally formed in the mutant embryos (Figure 5O-5R), suggesting another role of *pax1* genes for the aHM development in PP1 at a stage later than the PP1 formation.

In addition to RA and *pax1*, the membrane protein Alcam, which accumulates in the PP epithelium to stabilize the bilayered PP morphology [31], its accumulation was low in the PP1 and R2, but high in the C2 and more posterior PPs (Figure 6), suggesting that the R2-C2 border may also have separated the morphogenetic process of the endodermal epithelial cells. Taken together, we concluded that 2 distinct developmental processes proceeded in the pharyngeal endoderm either anterior or posterior to the R2-C2 border and that these processes were subsequently integrated to form the zebrafish PP2 (Figure 7Q). In other words, we clearly revealed that HA and BA1 were compartmentalized by distinct mechanisms and subsequently integrated by the dynamic endoderm of PP2. Since the emergence of R2, the caudal limit of HA, and C2, the anterior limit of BA1, are spatially separated in the zebrafish, we succeeded in dissecting each process, and in finding the obvious mechanistic interface of the PA development and the coordination process.

**Figure 6.**
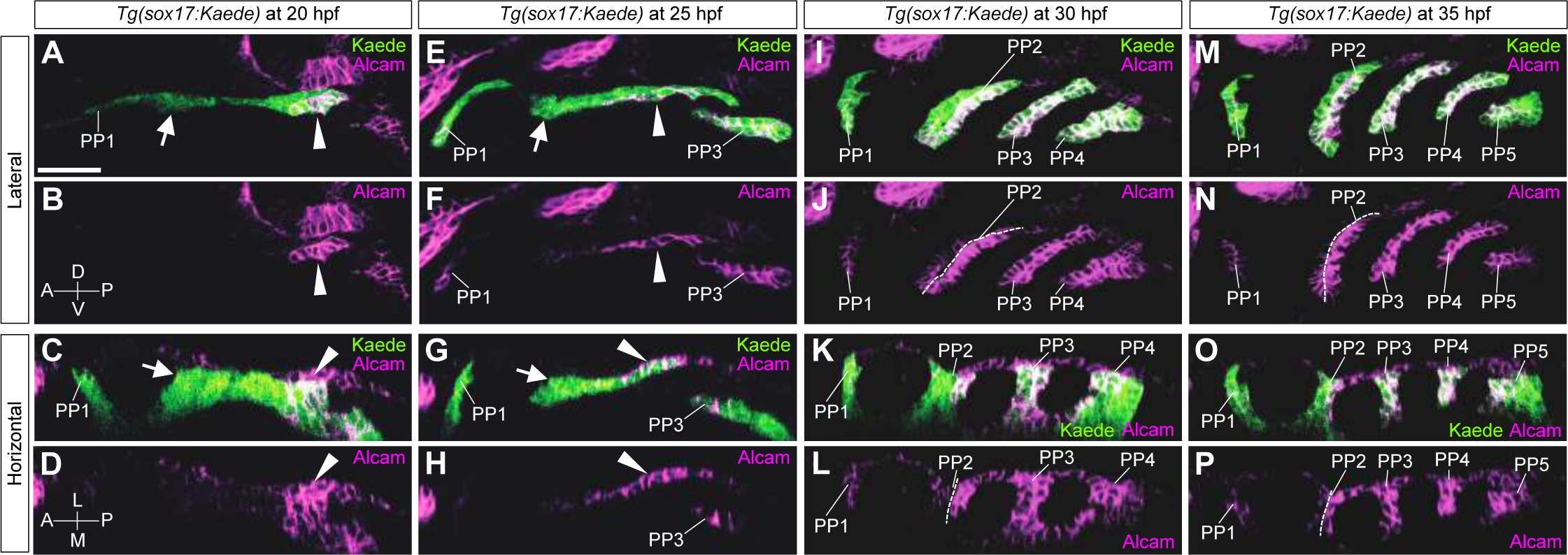
Expression analysis of Alcam in the PP endoderm of *Tg(sox17:Kaede)* (A-D) At 20 hpf, strong expression of Alcam was evident in C2 but hardly detected in PP1 and R2 endoderm. (E-H) At 25 hpf, Alcam was high in PP3 and the caudal aspect of PP2 but almost absent in PP1 and the rostral part of PP2. (I-L) At 30 hpf, high accumulation of Alcam was detected in PP3, PP4 and the caudal aspect of PP2 whereas it was very low level in PP1 and the rostral aspect of PP2. (M-P) At 35 hpf, high accumulation of Alcam was detected in PP3, PP4, PP5 and the caudal aspect of PP2 whereas it was very low level in PP1 and the rostral aspect of PP2. A, anterior; P, posterior; D, dorsal; V, ventral; L, lateral; M, medial; PP1–5, the first to fifth pharyngeal pouches; arrows, R2; arrowheads, C2. Scale bar, 50 μm.

**Figure 7.**
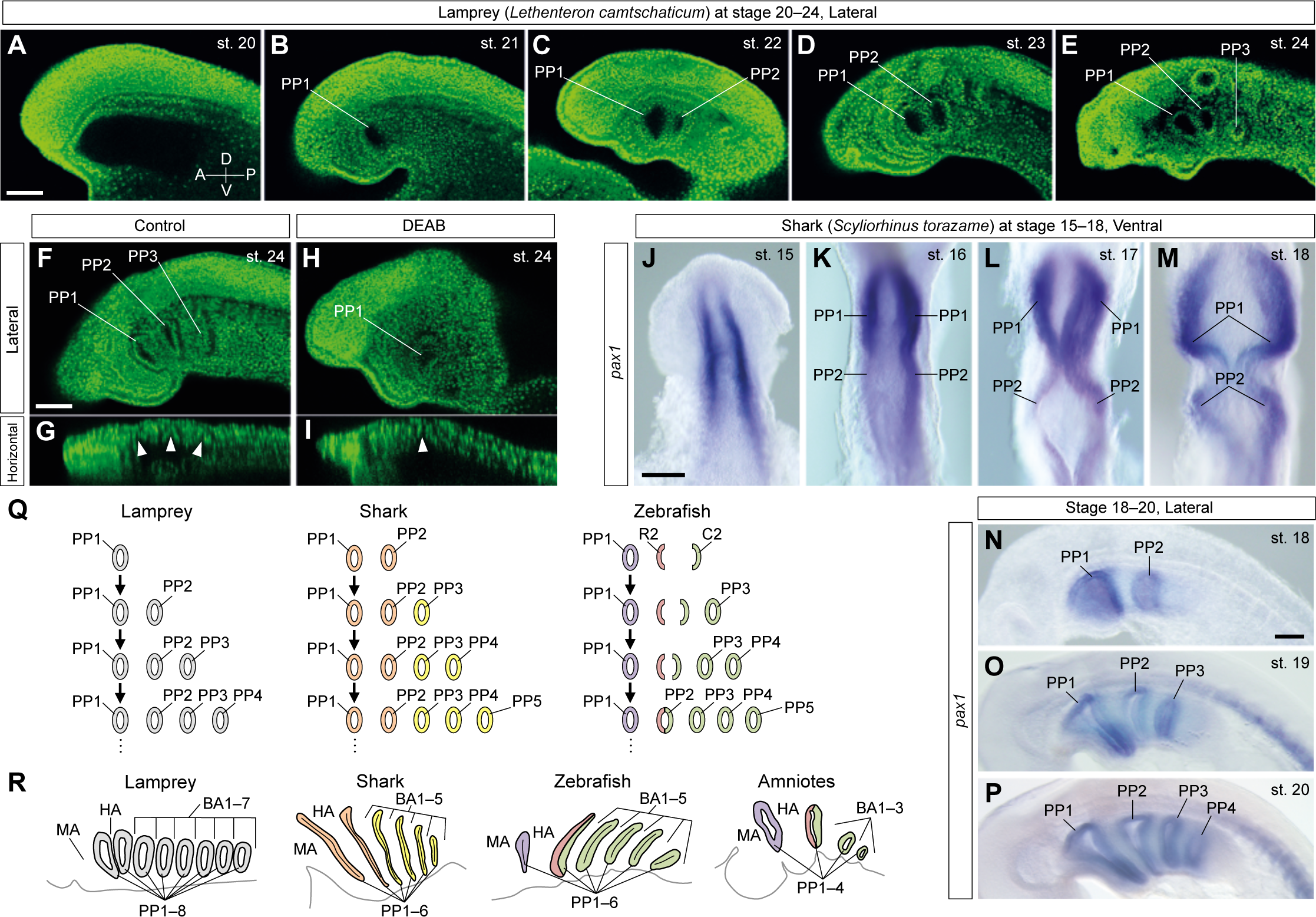
PP development in lamprey and shark embryos and the styles of the vertebrate PP segmentation. (A-E) PP development in the Japanese lamprey *(Lethenteron camtschaticum)* at stages (st.) 20–24. Only PP1 was formed at st. 21 (B), and PP2 appeared at st. 22 (C). Subsequently, PP3 was formed at st. 23–24 (D and E), showing that lamprey PPs, including PP2, were sequentially formed in an anterior to posterior order. (F-I) PP morphologies in control and DEAB-treated lamprey embryos at st. 24. In control embryos, PP1–3 were formed (F and G), whereas PP2 and PP3 were not formed in the DEAB-treated embryos (H and I). Arrowheads indicate the PP positions in optical horizontal sections (G and I). (J-P) PP development in the cloudy catshark (*S. torazame*) at st. 15–20. *In situ* signals from *pax1* probes visualized the lateral pharyngeal endoderm at st. 15, before PP segmentation was initiated (J). At st. 16, the endodermal epithelium formed 2 outpocketings per side (K). These pockets continued on to develop PP1 and PP2 (L and M), suggesting that the PP1 and PP2 formations occur simultaneously in shark development, as also seen in osteichthyans. After development of PP1 and PP2, PP3 and PP4 were sequentially formed at subsequent stages (N-P). *(Q* and *R)* Schematic summaries of the styles of PP development in the vertebrates. In zebrafish, PP1 (purple), R2 (red) and C2 (green) endoderm initially develop to segregate the regions of MA, HA and BAs, respectively. The dynamic process of the PP2 development subsequently integrates the anterior and posterior PAs to accomplish a theme-less series of PAs. In the shark, PP1 and PP2 (orange) are simultaneously generated to segregate the anterior PAs; whereas the posterior PPs (yellow) are sequentially formed. In the lamprey, however, all PPs (gray) are sequentially formed in an anterior to posterior order. The combinatorial styles of PP segmentation may be a conserved feature of gnathostomes (R). A, anterior; P, posterior; D, dorsal; V, ventral; BA1–7, the first to seventh branchial arches; HA, hyoid arch; MA, mandibular arch; PP1–8, the first to eighth pharyngeal pouches. Scale bars, 100 μm (A and F), 200 μm (J and N).

### PA Development Evolved in the Jawed-Vertebrate Lineage

When pharyngeal segmentation in vertebrates is compared with that in amphioxus and hemichordates, a temporal difference is found in the second segment of the endoderm: PP2 is formed simultaneously with PP1 in vertebrates, whereas the second gill slit is formed after the formation of the first gill slit in amphioxus and hemichordates [24, 27]. In order to gain insight into the evolutionary process of the pharyngeal segmentation in vertebrates, we investigated PP development in the lamprey, which lacks jaws and operculum. In contrast to those of osteichthyans, the anterior PPs during lamprey development were sequentially formed in an anterior to posterior sequence, as were the posterior PPs and the gill slits (Figure 7A-7E and 7Q). In addition, DEAB treatment impaired the PA development posterior to the PA2, including loss of the rostral aspect of PP2, in lamprey embryos (n=19/19, Figure 7F-7I). Considering that the HA segmentation by the rostral PP2 is resistant to RA deficiency in zebrafish and amniotes, lampreys appear to lack a developmental mechanism to establish HA without RA, resulting in the loss of HA in the treated embryos (Figure 7F-7I). Thus, the anterior PA development, especially the development of the rostral aspect of the PP2, was different between osteichthyans and lamprey, correlating with the huge morphological difference seen in the anterior PA derivatives between gnathostomes and cyclostomes.

This unexpected difference in PA development between lamprey and osteichthyans led us to conceive an evolutionary hypothesis that the developmental mechanism of PA was modified during the evolution of gnathostomes or, jawed vertebrates. To inquire whether osteichthyan-type development of the pharyngeal endoderm, in which the anterior 2 PPs are simultaneously formed, is conserved in gnathostomes, we next examined the PP development in a shark (*Scyliorhinus torazame)* of chondrichthyans, an extant sister group of osteichthyans [41]. The shark PP1 and PP2 simultaneously appeared as 2 pairs of broad lateral swellings in the *pax1* -positive endoderm, and these subsequently matured into the bi-layered PP morphology (Figure 7J-7M). Similar to those in osteichthyans, the more posterior PPs were sequentially formed in an anterior to posterior order (Figure 7N-7P). Although it remains elusive as to whether shark PP2 development is carried out by the same mechanism as in osteichthyans or not, the simultaneous emergence of PP1 and PP2 was clearly identical to that seen in the osteichthyans (Figure 7Q-7R).

## DISCUSSION

### Dissection of Dynamic Endoderm in Zebrafish PP2 Development Revealed Separate Development of Anterior and Posterior PAs and Their Integration

Generally, segmentation in animal development is carried out in a simultaneous manner, as seen in the *Drosophila* germ-band formation [42], or in a sequential manner, as represented by vertebrate somitogenesis[43], Interestingly, PA segmentation has been considered to be peculiarly achieved by the combined use of these distinct styles, although the developmental basis of their integration process has been entirely unknown. Zebrafish PA development, where dynamic epithelial remodeling of the PP-forming endoderm has been shown to take place, is a good model to resolve the elusive issue of development of the vertebrate head [31-34]. In this present study, by using precise live imaging in zebrafish, we found the 2 endoderm bulges of R2 and C2 in the future PP2 endoderm and uncovered their dynamic integration process forming PP2. Our cell-tracing experiments in the endoderm clearly revealed the direct contributions of R2 and C2 to the rostral and the caudal aspects of PP2, respectively. Their rostrocaudal identities in the PP were separately determined at an early timing in the PP segmentation, as evidenced by the gene expression and cell-ablation experiments. Importantly, the intermediate endodermal cells between R2 and C2 did not contribute to the formation of the PA-derived skeletons, although they were required for a tight arrangement of the anterior and posterior PAs. These results suggest that the anterior and posterior PAs were independently formed by distinct endodermal development (Figure 7Q). Subsequently, these distinct domains became integrated to form the systematic PA-derived organs by the dynamic epithelial transformation causing maturation of PP2 in the HA-BA border (Figure 7R).

Based on our results, we can adequately propose a novel view for vertebrate PA development; that is, HA and BAs are independently established by distinct developmental mechanisms for R2 and C2 endoderm, respectively. Given our viewpoint, the posterior PA-specific defects and the PP2 insufficiency, which are commonly reported as being phenotypes of RA-deficient vertebrates [17-20], are explained more reasonably as defects brought about posterior to C2. Similarly, the *pax1* mutants of teleost fish [21] and the *Ripply3* mutant mouse [22] show the anterior limits of the PP defects in the C2 endoderm. We previously proposed that developmental system drift should have occurred in the posterior PA segmentation among fish and mouse [21]. Notably, the anterior limit of the pharyngeal endoderm, where that drift has occurred, is also the C2 endoderm, strongly suggesting the conserved developmental interface between HA and BAs or the developmental independency of R2 from the posterior PAs in gnathostomes. Although *tbx1* is required not only for posterior PA development but also for HA development [44], our idea is not controversial because the expression of *tbx1* is independent from *pax1* function in HA endoderm; whereas that in BA endoderm is a *pax1* -dependent one [21]. This difference suggests that there are distinct gene regulatory networks between anterior and posterior PAs, rather highlighting the developmental independency of HA from BAs.

In addition to the separate establishment of HA and BA1, our study also revealed their integration process. Interestingly, the endodermal cells between R2 and C2 contributed to a part of PP2; however, they were not necessary to develop the HA- and BA1-derived skeletal elements, suggesting that they functioned developmentally like a glue to bind 2 PAs. This integration process or the endodermal cells identical to the intermediate endoderm have not been found in other species. Thus, we also speculate that the PA development largely depends on the endodermal epithelial morphogenesis, which is divergent among vertebrates. Further investigation by live-imaging analysis and cell-tracing experiments for endodermal behaviors of other vertebrates, especially amniotes, is required to shed light on whether the integrative development of PAs by the dynamic endoderm is common in vertebrates or not.

### Evolution of Pharyngeal Segmentation Possibly Contributed to Drastic Innovations of PA Derivatives

Whereas the vertebrate oropharyngeal regions display a huge variety of adaptive morphologies, the vertebrate PAs are fundamentally conserved in their architecture, developmental overview, and gene expression patterns[4]. Even in amphioxus and hemichordates, which diverged before the emergence of the vertebrates, the endodermal segmentation and expression of pharyngeal genes are largely conserved as in the vertebrate PPs [4, 23-26]. In the light of this information, our finding of the differential regulation of PP development between gnathostome species and the lamprey is surprising. Not only the RA dependence but also the timing of lamprey PP development was different from those of gnathostomes (Figure 7Q). The sequential segmentation of all pharyngeal segments in the lamprey was rather similar to that of the endodermal gill slits in the non-vertebrate deuterostomes. Additionally, the simultaneous segregation of the anterior PAs in shark embryos was identical to that in osteichthyans (Figure 7Q), indicating that the ancestral style of the PP development seen in the lamprey had been modified in the gnathostome lineage. Strikingly, this modification took place in the endoderm corresponding to R2, which we identified in the present study (Figure 7R).

What is the significance of the R2 acquisition for the morphological evolution of the vertebrate head? We suppose that the R2 acquisition probably contributed to the evolution of the opercular system, which is conserved in osteichthyans [39]. A recent study on a fossilized placoderm, *Entelognathus*, which has the opercular and branchiostegal rays, suggests that the osteichthyan-like pharyngeal system exists in the stem gnathostomes [45]. Furthermore, it has been also suggested that the chondrichthyan affinity of acanthodians, which possess a hyoidean gill cover with branchiostegal rays, implies unique evolution of the chondrichthyan pharyngeal system composed of septal gills [41, 46, 47]. Although the skeletal elements of the operculum have been lost during tetrapod evolution, the embryonic opercular flap, which is derived from the Shh-expressing HA, encloses the posterior pharyngeal region during amniote development [4, 39]. We identified the R2 endoderm, which directly contributed to the operculum, including the Shh-expressing cells in the zebrafish HA. Thus, we propose that the HA development distinct from that of the BAs should have been acquired in the stem gnathostomes, being the crucial basis for the novel pharyngeal system of the hyoidean operculum leading to the extent osteichthyans (Figure 8).

**Figure 8.**
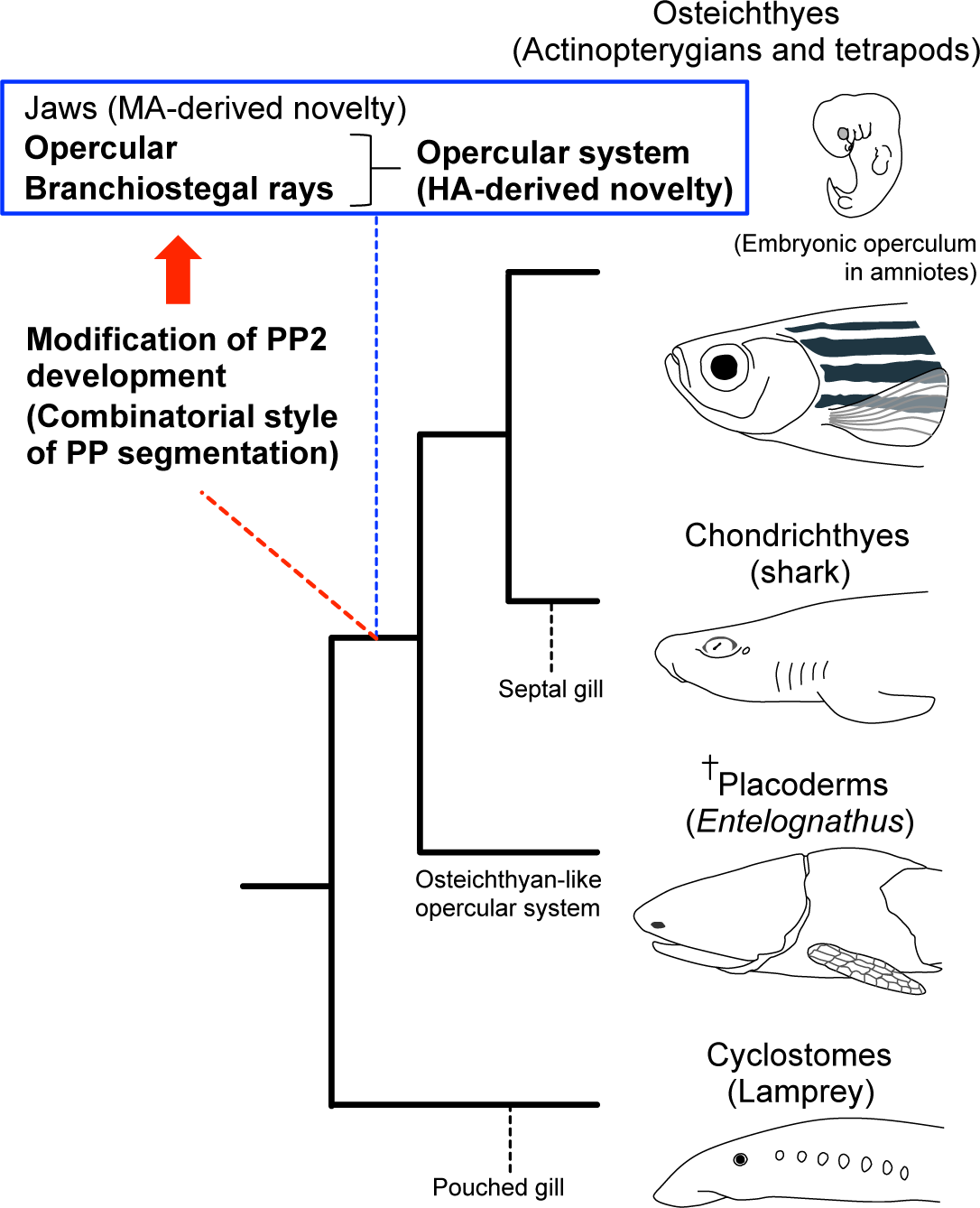
Evolution of PP development and pharyngeal systems in vertebrates. The development of PP2 was modified, and R2 region was acquired in the common ancestor of gnathostomes, resulting in the developmental separation of the anterior PAs from posterior PAs. This modification might significantly contribute to the evolution of the hyoidean operculum in gnathostome lineage, leading to the extent osteichthyans.

It is still unclear how the development of PP1 and the rostral layer of PP2 are regulated. The independency of the PP1 development from other PPs has been indicated by the results of previous studies on zebrafish [16, 29, 48] and mouse development [44, 49]. We showed that the lamprey PP1 was formed independent of RA signaling. Additionally, in amphioxus, the formation of the first gill slit is less affected by *pax1/9* knockdown than that of the other gill slits, which exhibit severe defects [50]. These findings imply the evolutionally conserved independency of the first endodermal bulges from the others. Therefore, the development of the vertebrate PPs, especially in gnathostomes, can be possibly considered as involving tripartite sections, i.e. PP1, the rostral layer of PP2, and more posterior region (Figure 7R). Significantly, these endodermal sections correspond to the interfaces of 3 streams of the cranial NCCs composing the MA, HA, and BAs. Therefore, we propose the possibility that the modifications of the endodermal segmentation reinforced the topological restrictions of the NCC streams in the PAs. Further studies on the development of PPs may answer one of the biggest issues regarding development of the vertebrate head, that is, the logic for the coordination between pre-patterned NCCs and endodermal segmentation.

## METHODS

### Zebrafish, shark, and lamprey embryos

Zebrafish with the TL2 background were used as the wild type, as described previously [51]. Collected embryos were incubated at 28°C. Embryos that would be fixed later than 25 hpf were treated with 0.003% 1-Phenyl-2-thiourea (PTU) from 10 hpf until fixation to inhibit melanin synthesis. Eggs of the cloudy cat shark *(Scyliorhinus torazame)*, which were collected from the established breeding colony in the Hekinan Seaside Aquarium, were grown at 16°C; and their developmental stages were determined according to the morphological criteria of a related species *(S. canícula)* [52]. Embryos of lampreys *(Lethenteron camtschaticum)* were collected as described previously [53], and their developmental stages were determined as described earlier [54]. This study was performed in accordance with the Guidelines for Animal Experimentation of National Institutes of Natural Sciences, with approval of the Institutional Animal Care and Use Committee (IACAC) of the National Institutes of Natural Sciences.

### Transgenic zebrafish and mutagenesis

*Tg(sox17:EGFP)* and *Tg(RARE:Venus)* were used in this study. For generation of *Tg(sox17:Kaede)*, the Kaede cDNA fragment from pKaede-S1 (MBL) and the same promoter sequence as used for the *Tg(sox17:EGFP)* were combined and cloned into the pSK-tol2B vector [55]. Transgenesis was performed by using the Tol2 system [56]. For CRISPR/Cas9-mediated knockout, target sequences were determined by using the ZiFiT Targeter [57]. Construction of guide RNA vectors and preparation of sgRNA and Cas9 mRNA were performed as described in previous reports [58, 59]. The mutation efficiency was assessed by performing a T7 endonuclease assay [60] using the following primers: 5’-TTG ATT TAG GTC ATG TGT GTT ATA TG-3’, 5’-TTT GTT TGT AGT CCC GTA TGT TTT T-3’ for *pax1a* and 5’-GTT TTT CTG ACA ATG CAA AAA GTG-3’, 5’-CGT ATT TCC CAA GCA AAT ATC C-3’ for *pax1b*. Details of *pax1a* and *pax1b* mutagenesis and sequences of sgRNAs are described in the Supplementary information (Fig. S3). For microinjections, 1 nl of each injection solution (Tol2 transgenesis: 25 ng/μl plasmid, 50 ng/μl Tol2 mRNA, 0.2M KCl, and 05% phenol red; CRISPR/Cas9: 25 ng/μl sgRNA, 100 ng/μl Cas9 mRNA, 0.2M KCl, and 0.05% phenol red) were injected into one-cell stage zebrafish embryos by using an IM300 micro injector (Narishige).

### Imaging

Living embryos of *Tg(sox17:EGFP)*, whose chorions had been manually removed, were anesthetized with 0.02% ethyl-3-aminobenzoate methanesulfonate (MS-222) in 1/3 zebrafish Ringer. For observations with a Leica SP8, the embryos were moved to mounting medium (0.15% low-melting-point agarose, 0.02% MS-222, 0.003% PTU in 1/3 zebrafish Ringer) and individually set in the medium on a glass bottom dish. Positions of the embryos were manually turned by a tungsten needle with an eyelash on its tip. Z-stack images were taken at 10-minute intervals, and the stack images were processed with an LAS X (Leica) to make 3D images, optical sections, and movies. For imaging with a Zeiss Lightsheet Z.1, the anesthetized embryos were mounted as previously described [61]. Images taken at 10-minute intervals were processed with ZEN Black (Zeiss) and subsequently with Imaris (Bitplane) to make movies.

### Photoconversion and cell ablation

Embryos of *Tg(sox17:Kaede)* for photoconversion or *Tg(sox17:EGFP)* for cell ablation were mounted as described above. Photoconversion was performed with a Leica SP8 using a 405-nm diode laser. Regions of interest (ROI) in the Kaede-expressing endoderm at 20 hpf were converted by using the ROI tool in LAS X (Leica). Converted embryos were released from the gel and incubated at 28°C, and were observed again at 48 hpf. Cell ablations with an IR-LEGO system were performed as previously described [38]. A high-power flash irradiation from an IR laser (80 mW for 8 ms) was performed a few times until the EGFP signals of target regions had been eliminated. Operated embryos were released from the gel and incubated at 28°C until subsequent experiments could be performed. For the control experiment for cell ablation, *Tg(sox17:EGFP)* embryos were injected with mRNA of histone H2A-mCherry at the one-cell stage to visualize cell nucleus in the live condition. At 20 hpf, embryos were scanned with a Nikon A1 before ablation and moved to the IR-LEGO. After infrared irradiation on the IR-LEGO, embryos were immediately moved to the Nikon A1 and scanned again to evaluate off-target damage to cells of adjacent PAs. This procedure was repeated, and the operated embryos were fixed in 4% PFA/PBST and stored in methanol at -20°C for *in situ* hybridization with a *dlx2a* probe and for immunohistochemistry with anti-GFP antibody to assess CNCCs in the PA.

### Staining

Whole-mount *in situ* hybridization of zebrafish was performed as described previously [62]. For double-fluorescence *in situ* hybridization experiments, anti-DIG-POD (Roche) and anti-FITC-POD (Roche) were used to detect each hapten in RNA probes. Fluorescent signals were detected with a TSA Plus Cy3/fluorescein system (PerkinElmer). Plasmids for probes of *dlx2a, nkx2.3*, and *shha* [63] were kindly donated by Drs. M. Hibi, Y. Kikuchi and S. Krauss, respectively. Primers for cloning other probes of zebrafish genes were as follow: 5’-ATG CTT TCG TGT TTT GCA GAG CAA ACA TAC-3’, 5’-TTA CGA GGA TGA GGT AGA AAG GCT GAG TCC-3’ for *paxla*; 5’-ATG CAA ATG GAT CAG ACG TAC GGG GAG GTG-3’, 5’-TTA TGA GTC TGA GAG TCC ATG AAC AGC GCT-3’ for *pax1b*; 5’-ATG ATT TCA GCA ATA TCA AGC CCG TGG CTG-3’, 5’-TTA TCT GGG TCC GTA GTC ATA ATT AGT CGG-3’ for *tbx1;* and 5’-AAA TCT CAC GAT AGG CTC CCT G-3’, 5’-AAA GTA CTC CTG ATT GCA GT-3’ for *fgf3*. Immunostaining was conducted as previously reported [64] by using primary antibodies anti-Alcam (1:500; Developmental Studies Hybridoma Bank, University of Iowa City), anti-Kaede (1:400; MBL, PM012), anti-GFP (1:400; Abcam, AB13970), followed by Alexa Fluor-conjugated secondary antibodies (Invitrogen). Bone and cartilage staining of zebrafish larvae was performed as previously described [65]. Whole-mount *in situ* hybridization of shark embryos was performed as per the zebrafish method. *S. torazame Pax1* was cloned by use of the following primers: 5’-ATG GAT CAG ACT TAC GGA GAG GTT AAG G-3’, 5’-TTA CGT ACT GGA GGC CGG GAT TG-3’. For nuclear staining of lamprey embryos, dechorionized embryos were fixed in 4% PFA/PBST and stored in methanol at -20°C until used. Rehydrated embryos were treated with RNaseA, and subsequently stained overnight at room temperature with YOYO1-Iodide in PBST (1:2000; Thermo Fisher Scientific). The stained embryos were rinsed with PBST several times, dehydrated with methanol, and soaked in BABB (benzyl alcohol/ benzyl benzoate, 1:2 ratio) prior to confocal imaging.

### DEAB treatment

DEAB (N,N-diethylaminobenzaldehyde) stock, which was stored at -20°C, was prepared at a 100 mM concentration in DMSO. DEAB treatment was conducted at final concentrations of 10^−4^M for both zebrafish (from 10 to 20 hpf) and lamprey (from st.16 to st.24), as previously described [17, 66]. The embryos were cultured in a dark incubator, at 28°C for zebrafish and 16°C for lampreys. Because of long-term development of lamprey embryos, the DEAB solution bathing these embryos was replaced with fresh solution once per day. Control embryos were treated under the same conditions, but with 0.1% DMSO only.

## Acknowledgements

We thank Dr. Y. Kikuchi for *Tg(sox17:EGFP)* zebrafish, a plasmid of the *sox17* promoter, and *nkx2.3* probe; Dr. S. Higashijima for technical support in CRISPR/Cas9 introduction; Dr. S. Krauss for the *shha* probe; Mr. M. Masuda and staff at the Hekinan Seaside Aquarium for collecting *S. torazame* embryos; Drs. S. Kuraku and K. Onimaru for shark information; Ms. H. Utsumi for technical support; Mrs. K. Takashiro for zebrafish maintenance; the Spectrography and Bioimaging Facility, NIBB Core Research Facilities, for technical support; Mmes. S. Ukai and R. Nobata for assistant secretary support; and all members of the S.T. laboratory for helpful discussions. This work was supported by the program Grants-in-Aid of the Japan Society for Scientific Research on Innovative Areas to S.T. [no. 24111002] and by an NIBB Young Researcher Fellowship (2016) to K.O.

## Author contributions

K.O. and S.T. conceived of and designed the research; K.O. performed all the experiments; K.O. confirmed and analyzed all the data; H.W. gathered the lamprey materials; K.O., S.T., and H.W. wrote and edited the manuscript.

